# Characterization of spatiotemporal dynamics in EEG data during picture naming with optical flow patterns

**DOI:** 10.1101/2022.11.24.517789

**Authors:** V. Volpert, B. Xu, A. Tchechmedjiev, S. Harispe, A. Aksenov, Q. Mesnildrey, A. Beuter

**Affiliations:** Institut Camille Jordan, UMR 5208 CNRS, University Lyon 1, 69622 Villeurbanne, France; EuroMov Digital Health in Motion, Univ Montpellier, IMT Mines Ales, Ales, France; CorStim SAS, Montpellier, France

**Keywords:** human EEG data, spatiotemporal patterns, optical flow methods, picture-naming task

## Abstract

We present an analysis of the spatiotemporal dynamics of the oscillations in the electric potential that arises from neural activity. Depending on the frequency and phase of oscillations, these dynamics can be characterized as standing waves or as out-of-phase and modulated waves, which represent a combination of standing and moving waves. We characterize these dynamics as optical flow patterns, in terms of sources, sinks, spirals and saddles. Analytical and numerical solutions are compared with real EEG data acquired during a picture-naming task. Analytical approximation of standing waves allows us to establish some properties of pattern location and number. Namely, sources and sinks have mainly the same location, while saddles are located between them. The number of saddles correlates with the sum of all the other patterns. These properties are confirmed in both the simulated and real EEG data.

## 1 Introduction

Developing adequate theoretical frameworks to describe the extreme complexity of the spatiotemporal electrical dynamics of 3D cortical neural tissue in electroencephalogram (EEG) recordings still remains a real challenge. These frameworks have a key role in the way we understand how cortical electrical activity functionally contributes to the dynamics in human behavior.

EEG records the electric potentials in the brain at the scalp. The amplitude and phase of the oscillations that characterize the dynamics of electric potentials depend on the spatial locations of the electrodes. Spatiotemporal dynamics in EEG data has been studied since the 1930s (see [1] and the references therein) when oscillations in the electric potentials in the brain were found to originate from specific brain sources (originally called *focuses*) located in the occipital lobes. It was suggested that these sources can shift within limited areas in the brain, giving rise to phase and amplitude shifts in the signals recorded at different electrodes, and interpreted as moving waves, also referred to as *travelling waves.* The contemporary use of the term *travelling waves* has come to be mathematically more defined and is a function which depends on the combination of variables *x* — *ct*, where c is the wave speed. Simple definitions consider travelling wave to have constant speed and amplitude, but general definitions acknowledge that speed and amplitude can be variable [2].

The phase and amplitude of these travelling waves have been extensively studied for different brain states in humans and in animals (see literature reviews [3,4]). Travelling alpha waves were found in [5] across four occipital-parietal electrodes during a visual cognitive interference task in human subjects. Periodic travelling waves along the frontal-occipital axis were also found during a cognitive control task [6] and suggested as a way of viewing slow sleep waves [7]. They were recorded during 15-20% of the observation time and their direction changed, being slightly more frequent for frontal-occipital waves pre-stimulus and occipital-frontal waves post-stimulus. Bidirectional travelling waves were found in [8] with posterior-to-anterior travelling waves being more frequent during visual input and anterior-to-posterior during rest. Travelling waves have been observed in the primary visual cortex where they were reduced when a wide part of the visual field is strongly stimulated [9]. They can also help in understanding language processing [10], such as that involved in semantic feature during lexical access [11]. It should be noted that the phase component of the signal observed in individual trials can be lost in across-trial average [12]. Moreover, oscillations are not simply plane waves but can be rotating like in sleep spindles [13,14] or spiral waves [15].

Approaches in analyzing brain dynamics, however, have focused on trial and group averages, as is the case with analyses of event-related potentials (ERP) using global field power (GFP), which has been associated with brain micro-states [16–20], brain sources, and networks for various cognitive tasks [21–23]. For example, GFP was used to compare the dynamics of phonological encoding between stroke patients and healthy subjects in [24]. Different approaches to analyze brain dynamics at scalp, sources and networks during picture naming task are discussed in [25]. Spatiotemporal dynamics of electric potentials have been characterized using block-matching motion estimation [26] and calculating peak amplitude trajectories [27] from topographic maps.

To characterize neural oscillation dynamics in EEG recordings that might be relevant to cognitive processes but lost in averaged data, we propose an individual- and trial-by-trial based approach inspired by optical flow methods used in computational vision models. With this approach, we determined types of spatiotemporal regimes (optical flow patterns, OFP) in simulated data of neural activity under alternate current stimulation as well as real EEG data recorded in healthy human subjects during a picture-naming task. This approach can be used to characterize any sufficiently smooth function *F*(*x, y, t*) that varies in space (*x, y*) and time *t*. Trajectories of points in space (*x*(*t*), *y*(*t*)) determined by this function constitute a vector field in a plane which can have singular points (nodes, focuses, saddles) characterizing the function *F*. This method can be applied to either the amplitude or phase of EEG signals treated as a (discrete) function of space and time.

This method has been used in [28] to analyze local field potentials (LFPs) measured in the visual cortex of anesthetized marmoset monkeys in the delta frequency range. Analysis of the phase fields revealed sources (unstable nodes), sinks (stable nodes), spirals (focuses) and saddles. When plane waves are present, singular points are absent, implying that plane waves and other patterns are mutually exclusive. Plane waves were found to be the dominating pattern 60% of the time, while spatial patterns were present in 20.4% of the time, and synchronized EEG (no spatial distribution) in 19.6%. Transitions between simple waves (synchrony, plane) were also less frequent than transitions from simple to complex waves. Complex patterns arise around preferential locations, as has been found in [29] and in local complex wave patterns in the phase velocity field of spontaneous dorsal brain activity in anesthetized mice [30]. It was observed that sources, sinks and saddles frequently coexisted while global plane waves inversely correlated with these patterns. Large-scale waves propagate preferentially in the anteroposterior direction, and the change of their direction was related to the emergence of sinks or sources. Location preferences of these patterns appear to be anatomically motivated, as suggested by the localized propagation in limited visual cortex subregions at rest [30] to wider propagation beyond the visual cortices during visual stimulation [31].

Depending on the frequency and phase as well as source location, we observed different types of dynamics in simulated spatiotemporal regimes (Section 2) that can be characterized as standing waves, out-of-phase standing waves, and modulated waves. The dynamics in the EEG data are qualitatively similar.

In Section 3, we characterize simulated dynamics with OFP and determine some properties of these patterns, such as their mutual location and number. These properties are then verified on real EEG data and additional general OFP found in the EEG data are described.

Finally, more specific properties of optical flow patterns evoked during the picture-naming task are described in Section 4.

## 2 Brain sources and spatiotemporal dynamics

### 2.1 Spatiotemporal dynamics in simulated data

In this section we present the results of numerical simulations with 3D realistic brain geometry using software SimNIBS [32]. This tool allows modelling of transcranial direct current stimulation (tDCS) with stimulating electrodes located at the brain surface. The model uses Poisson equation [33]. Since this equation is linear, it is thus possible to use it for modelling of transcranial alternating current stimulation (tACS) with several simultaneously acting time-dependent sources [32]. Numerical implementation is presented in Supplementary Materials B.

In numerical simulations, we use 30 electrodes including 3 tACS electrodes and 27 registering electrodes. We consider two stimulating electrodes and one return electrode, where stimulation observes the conservation of charges, that is, the sum of injected currents is equal to zero at any given time point. With this configuration, different regimes can be identified depending on frequency and phase at the two stimulating electrodes, and can be basically considered as three cases: 1) equal frequencies and phases, 2) equal frequencies and different phases, and 3) different frequencies and phases.

For the case of equal frequencies and phases, the dynamics corresponds to standing waves (Figure 1, left). All signals have constant amplitude and vanish at the same time points. The spatial 2D projection of the 30 point-wise simulated values on the circular domain is shown in Figure 2. Time oscillations in this case are synchronized; that is, spatial locations of maximums, minimums, and zeros do not depend on time.

**Figure 1.**
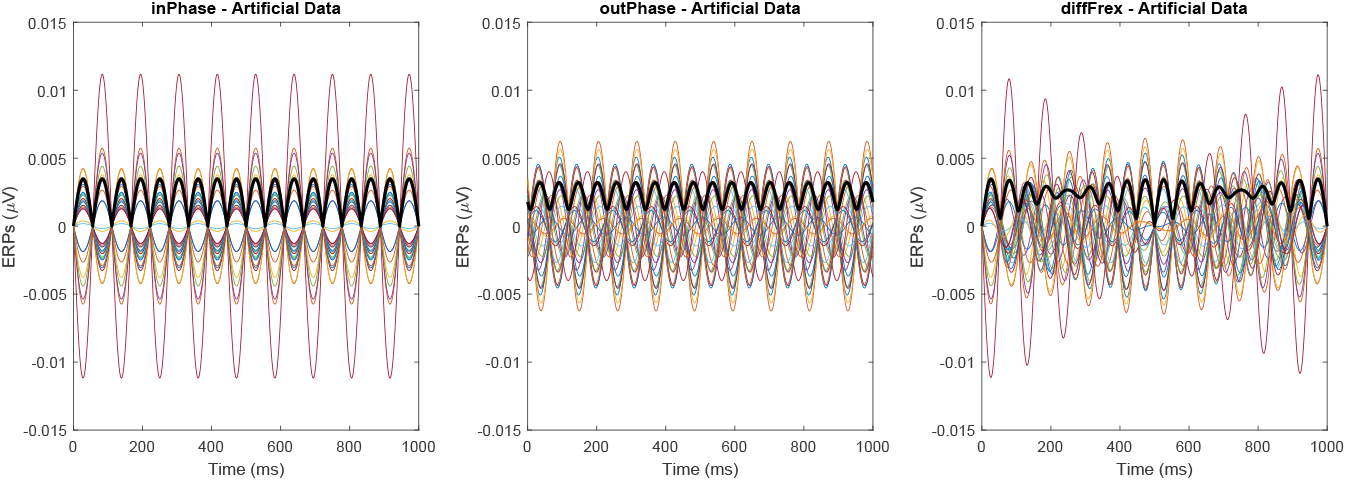
Simulations of tACS with three electrodes generated by the software SimNIBS. Each colored curve is the simulated electric potential time series at one of the 27 electrodes. The three heuristic cases are shown: 1) equal frequencies and phases (left), which results in standing waves; 2) equal frequencies and different phases (middle), which results in out-of-phase waves; and 3) different frequencies and phases (right), which gives rise to either out-of-phase standing waves or amplitude-modulated out-of-phase waves. Bold black lines showtime-dependent global field power (GFP).

**Figure 2.**
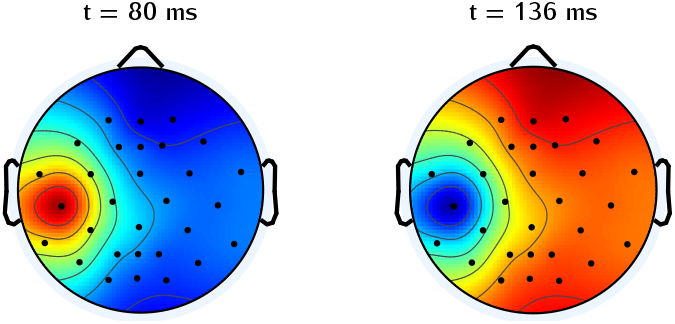
Spatial 2D projection (topographic map) of the 30 simulated electrodes for the case of equal frequencies and phases. The numerical solution is a standing wave with synchronized time oscillations. The two topographic maps shown are two snapshots of the solution with opposite values.

In the case of different phases, we obtain out-of-phase standing waves characterized by phase-shifted signals with constant amplitude (Figure 1, middle). If frequencies and phases are different, then the dynamics corresponds to an out-of-phase modulated wave (Figure 1, right) with signals periodically changing in amplitude and phase.

Figure 3 represents topographic maps of signal amplitude in consecutive time points during one period in the case of different frequencies. These topographic maps repeat for several periods and then change rotational direction. The upper and lower panels in this figure show similar distribution patterns of electric potential amplitude but of opposite sign (indicated by inverted hot/warm colors). One more property of this solution is that the rotation is not spatially uniform. Rotation is driven by an alternation of when forward and backward wave fronts propagate. One wave front propagates while the other is fixed, and then the other front propagates in apparent rotating motion while the front that just rotated remains stationary. In the solutions to this case, the maxima alternate between spatially displacing at approximately a constant speed, characteristic for travelling waves, and jumping to distant locations, characteristic of standing waves.

**Figure 3.**
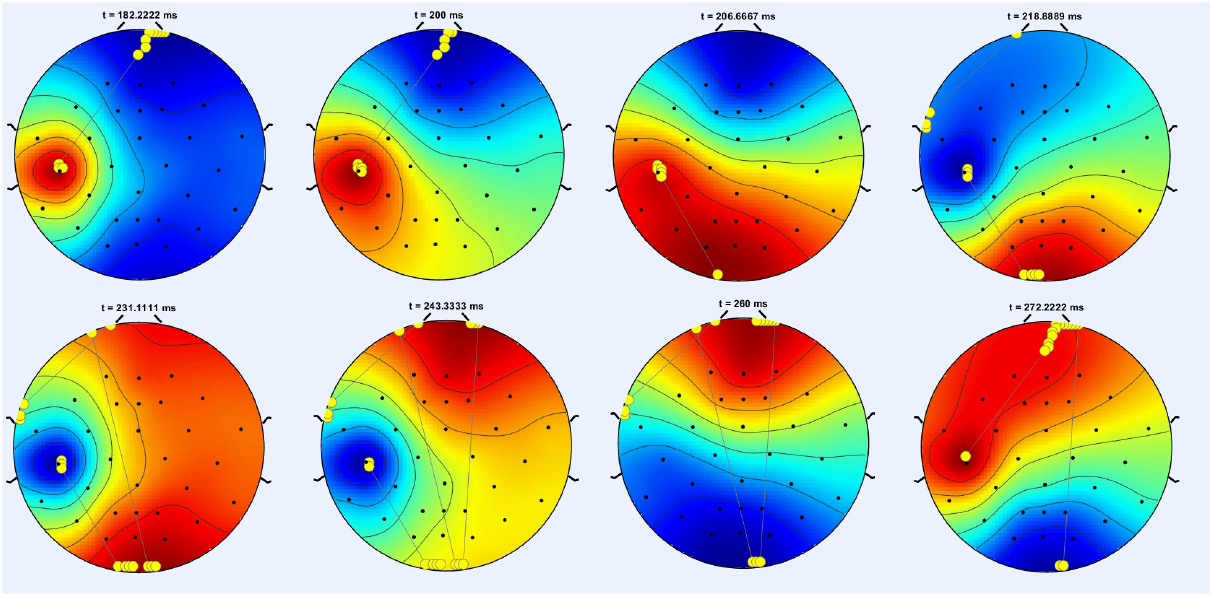
One period of a rotating wave in the case of different frequencies for the simulated data. Yellow dots show the maximum potential for a given time window corresponding to the given time point t shown and some time points prior to t. Straight black lines show the transitions of this maximum. Direction of rotation changes after several periods. Note that rotation is not uniform but corresponds to the fast transitions between brain states. This transition occurs through the propagation of the forward front followed by the propagation of backward fronts. Each map in the upper row is shown with a similar counterpart directly below it with the sign (hot/warm color) reversed.

Let us note that time-dependent GFP is periodic in the case of equal frequencies (Figure 1, left and center plots). The smallest average value is zero in the case of equal phases, and is positive for different phases. If frequencies are different, the average amplitude is not periodic.

The three regimes observed in the simulated data are qualitatively similar to the analytical solution (Supplementary Materials C). Standing waves are observed for equal frequencies and phases, out-of-phase standing waves for different phases, and modulated out-of-phase waves if frequencies are also different.

### 2.2 Spatiotemporal dynamics in EEG data

In this section we will consider spatiotemporal dynamics of the EEG data during a picture-naming task for 16 human subjects. Data collection and preprocessing are described in Supplementary Materials A. The analyses were performed on four frequency bands: delta (1–4Hz), theta (4–8Hz), alpha (8–13Hz), and beta (13–30Hz).

Like the analysis performed with simulated data, the real evoked EEG signal dynamics can also be described in terms of standing waves, out-of-phase standing waves and modulated waves.

An example of standing waves in EEG data during naming with 96 electrodes is presented in Figure 4–S17. Time 0 here corresponds to the onset of picture presentation. After the picture appears on the screen, the amplitude of the signal increases for approximately 300 ms, and then it drops back to baseline level. We can identify properties of standing waves during the first 300 ms. Recorded signals have sinusoidal oscillations and vanish at the same time points. The 2D spatial distribution of the electric potentials projected from 96 point-wise signals (electrodes) also has characteristics typical for standing waves, namely synchronized oscillations with fixed maxima, minima, and zero lines (Figure 4–S17).

**Figure 4.**
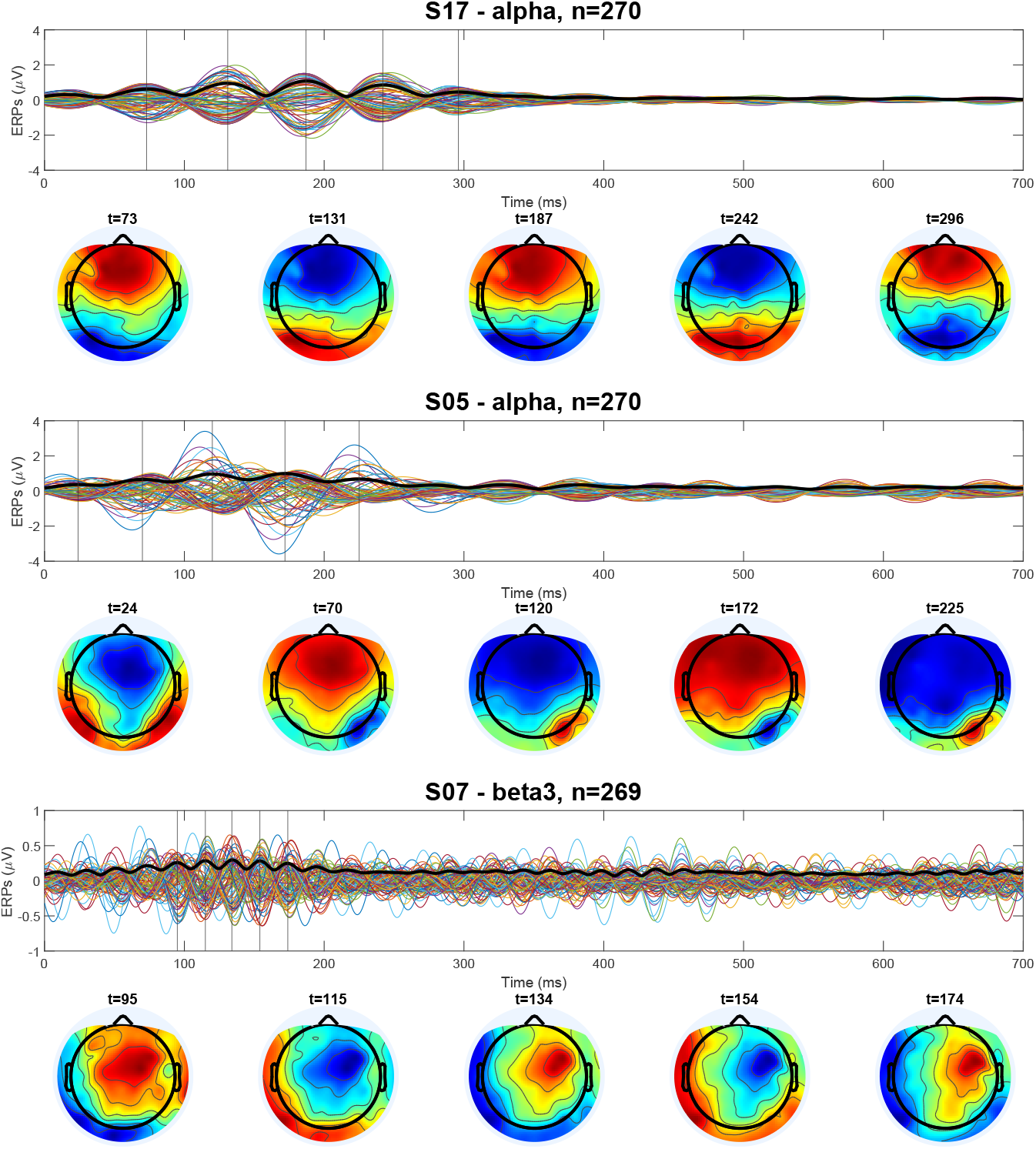
EEG signal time series averaged across all trials during naming (upper panes) in three healthy human subjects (S17, S05, S07), and the corresponding electrical potential topographic projection (topo plots) from 3D scalp coordinates (lower panes). Bold black curves in the signal plots represent the GFP. Vertical lines mark 5 maxima of the average amplitude correspond to the moments of time of topo plots below. An image appears on the screen at time 0. Image recognition and processing is associated with the ERP increase in the first 300 ms.

Out-of-phase standing waves were observed for one individual in the same frequency range (Figure 4–S05). Potential distribution in space is not exactly periodic in time (topographic plot/lower pane). Finally, an example of modulated out-of-phase waves for the *β* frequency range is shown in Figure 4–S07.

To summarize, in the analytical and numerical models with three stimulation sources, the spatiotemporal dynamics can be described by standing waves, out-of-phase standing waves and modulated waves. Spatiotemporal dynamics in the EEG data show similar behaviors. Moreover, the difference in the dynamics in Figure 4 (S17 and S05) can be related to equal or different frequencies at brain sources, as is the case for the analytical and numerical models. Therefore, a time-dependent Poisson equation with several sources should also be able to model the spatiotemporal dynamics of ERPs.

## 3 Optical flow patterns

In this section we will establish the connection between spatiotemporal dynamics described in the previous section and optical flow patterns. The definitions and the methods of analysis of these patterns can be found in [29, 30] (see also Supplementary Materials, C, D). We will formulate some hypotheses about the location and the number of patterns based on the analytical approximations. We will verify these hypotheses for the generated data and for the acquired EEG data.

### 3.1 Location and number of patterns for standing waves and other regimes

Analyzing spatiotemporal regimes (Supplementary Materials, C) allows us to make some hypothesis about location and number of patterns. In the case of standing waves, each maximum or minimum with increasing amplitude (in the absolute value) corresponds to a source (unstable node or focus) and with decreasing amplitude to a sink (stable node or focus). Since standing wave maxima and minima have fixed positions, and their amplitude is a periodic function of time, we have the following properties:

#### Property 1

*Sources and sinks of standing waves alternate occupying the same spatial locations. Their location does not depend on time.*

These properties will be verified below for all regimes and not only for standing waves. Furthermore, according to the properties of dynamical systems, sources (sinks) should be separated by saddles. Therefore, we will also analyze their mutual locations.

Some additional information about singular points can be obtained from the theory of dynamical systems. Let us consider a 2D vector field (*u*(*ξ, η*), *ν*(*ξ, η*)) in a closed manifold *S* without a boundary, such as a sphere in 3D space, (*ξ, η*) ∈ *S*. For each singular point (*ξ_i_*, *η_i_*), that is a point for which *u*(*ξ_i_,η_i_*) = *ν*(*ξ_i_,η_i_*) = 0, consider the following Jacobian

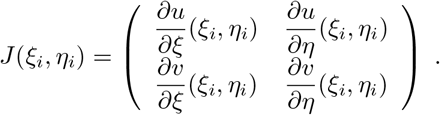

Assuming that the singular points are non-degenerate, det *J*(*ξ_i_, η_i_*) ≠ 0, *i* = 1,..., *n*, we can conclude that their number *n* is finite, and we can define the number

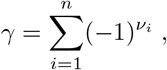

where *ν_i_* is the number of real positive eigenvalues of the matrix *J*(*ξ_i_,η_i_*). The number *γ* is called the topological degree or rotation of the vector field, and it is related to the winding number for plane vector fields.

Given that *ν_i_* = 0 for stable nodes (i.e., sink) and foci (spirals), *ν*_i_ = 2 for unstable nodes (sources), and *ν*_i_ = 1 for saddles, we thus have

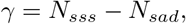

where *N_sss_* is the total number of sources, sinks and spirals, and *N_sad_* is the number of saddles.

We know that from dynamical systems, a vector field (*u_τ_*(*ξ,η*),*ν_τ_*(*ξ, η*)) continuously dependent on parameter *τ, γ* (*τ*) is in fact independent of *τ*. In terms of optical flow estimations which are a function of time, we therefore hypothesize

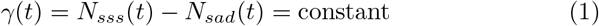

That is, the difference between the number of all singular points except saddles and the number of saddles does not depend on time, leading us to our second property.

#### Property 2

*The difference between the number of all patterns except saddles and the number of saddles does not depend on time.*

It is important to note that this property holds for the whole brain surface, and not only for standing waves but also for other regimes. If we consider only a part of the surface, as is the case of the EEG data, we should then consider the patterns crossing the boundary of the domain. If this information is not available, then this equality can be considered as an approximation and compared with the available data.

Another remark concerns the dependence of the location and number of patterns on the frequency band. In the case of standing waves, location and number are independent of the frequency. We will verify below whether this property is confirmed for both simulated and EEG data.

### 3.2 Optical flow patterns in simulated data

Depending on the phase and frequency in the simulated data, we observe standing waves, out-of-phase standing waves and modulated waves. In the case of standing waves, as from the earlier analysis, we can expect that sources and sinks have the same location in this ideal configuration. Furthermore, in analytical approximation, they coincide with the maxima and minima of the potential distribution. For the generated data, there is a number of stages of data processing which could influence the result. Let us recall that 30 signals were simulated as they would have been recorded at the scalp. These signals were then projected onto a circular domain, as is done for the real EEG data. However, the same signals were used to generate optical flow patterns with some other transformations (see Supplementary Materials, A). Different methods of data processing could possibly lead to some discrepancy in the results. We will verify Properties 1 and 2 for the simulated data.

#### Location of patterns

Figure 5 shows pattern position in time for the generated data. Since these data were generated with the two different frequencies 9Hz and 10Hz, patterns are only evaluated in the *alpha* frequency band (8 — 13Hz). In the first case with no mixtures (same-frequency, same-phase, upper panel), we observe only two locations of sources and sinks which overlap with each other^1^. Saddle patterns are alternated and located between source and sinks in a very regular way. The locations of these patterns are time-independent in this case.

For a more complex signal with mixed phases (middle panel) or mixed frequencies and mixed phases (lower panel), more patterns are generated and detected thereafter. Since the signal dynamics in these two cases are temporally and spatially (2D/3D) more complex, some variations are expected. However, we still observe very similar phenomena as in the first case: source/sink are most likely overlapped; saddles are found between source/sink. Their positions remain relatively stable – depending less on time.

#### Number of patterns

In the first case (the same frequency and phase), the numbers of sources and sinks are almost the same (17 and 18, respectively). There are approximately twice as many saddles (38). According to the theory of dynamical systems, two sources (sinks) are separated by a saddle point in such a way that the number of sources (sinks) and saddles is the same (plus/minus one, see Figure 5, upper panel). However, since sources and sinks replace each other periodically in time, while saddles are present all the time, then the time average number of saddles is twice as many than sources or sinks. In the second case (same frequency, different phases), after a noisy first 400-500ms, the number of patterns stabilize. Though the last case is most complex, the numbers of sources and sinks are similar while there are approximately twice as many saddles. All these observations agree with the hypotheses discussed in Section 3.1.

**Figure 5.**
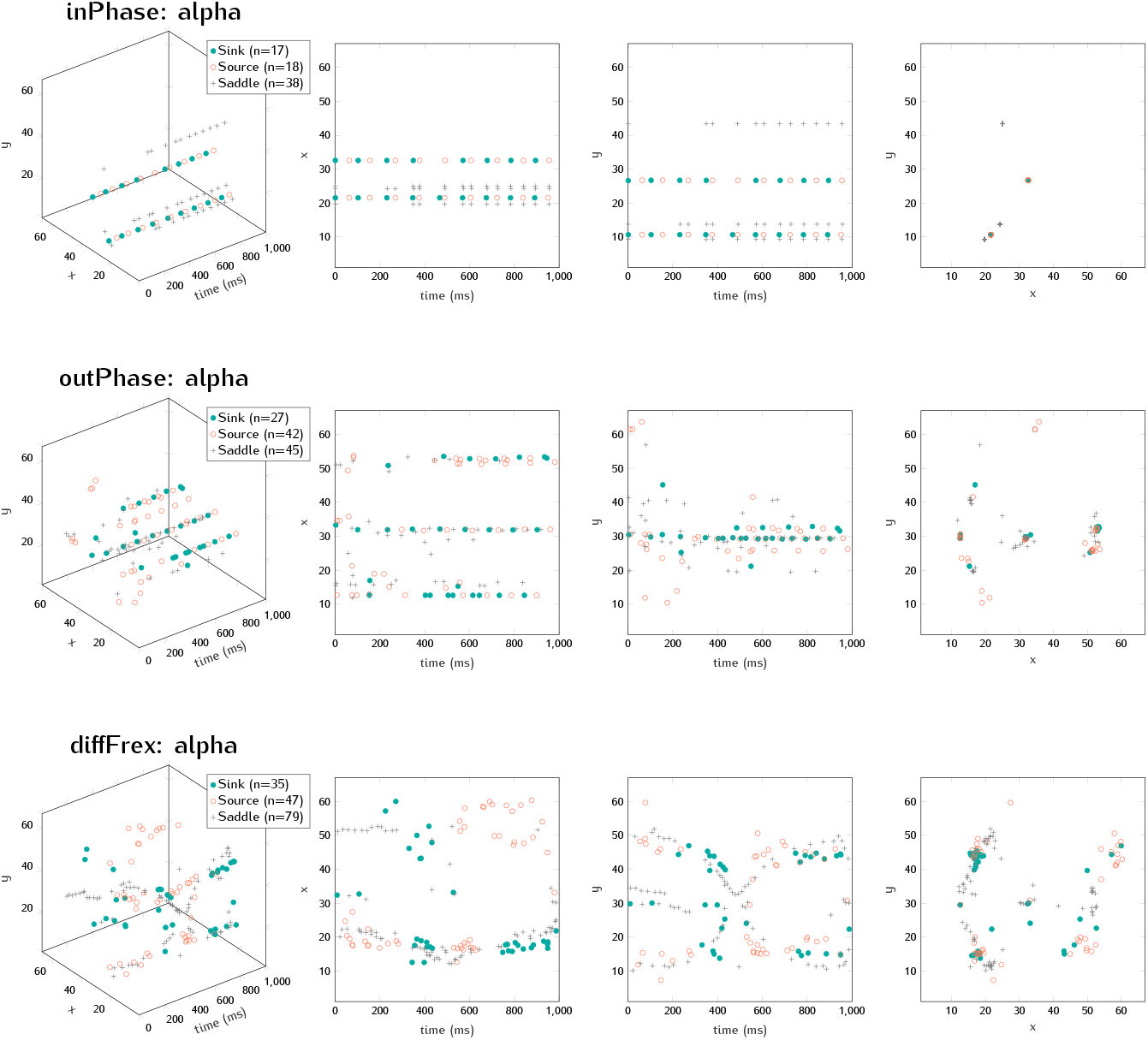
Time dependence of pattern locations for three simulated data: 1) inPhase – same frequency (9Hz from electrode positions AFz / POz), same phase; 2) outPhase – same frequency, different phases; and 3) diffFrex – different frequencies (9Hz from AFz and 10Hz from POz), different phases. Evaluated on amplitude and on *alpha* frequency band (8 — 13Hz). *n* is the number of patterns.

### 3.3 Optical flow patterns in EEG data

Results of Section 2 allow us to interpret EEG dynamics as standing waves, out-of-phase standing waves, and modulated waves. Analysis of optical flow patterns in Appendix D suggests that such regimes satisfy Properties 1 and 2. In the previous section, these properties were verified for the simulated data. We will now verify them for the real EEG data.

The EEG data for all 16 subjects and 270 trials are included in the analyses. The numbers of sources and sinks are approximately the same, similar to the number of spirals-in and spirals-out (Figure 6). Source/sink patterns occur about 3% more than spiral-type patterns. However, saddles have the largest share of the identified patterns – about 45% of the total number of patterns. Similar to the simulated data, the number of saddles was approximately equal to the total number of the other two patterns (sources and sinks). However, sources are partially replaced by spirals- out and sinks by spirals-in. We note that the analysis in Supplementary Materials Appendix D does not distinguish between sources (sinks) and spirals.

**Figure 6.**
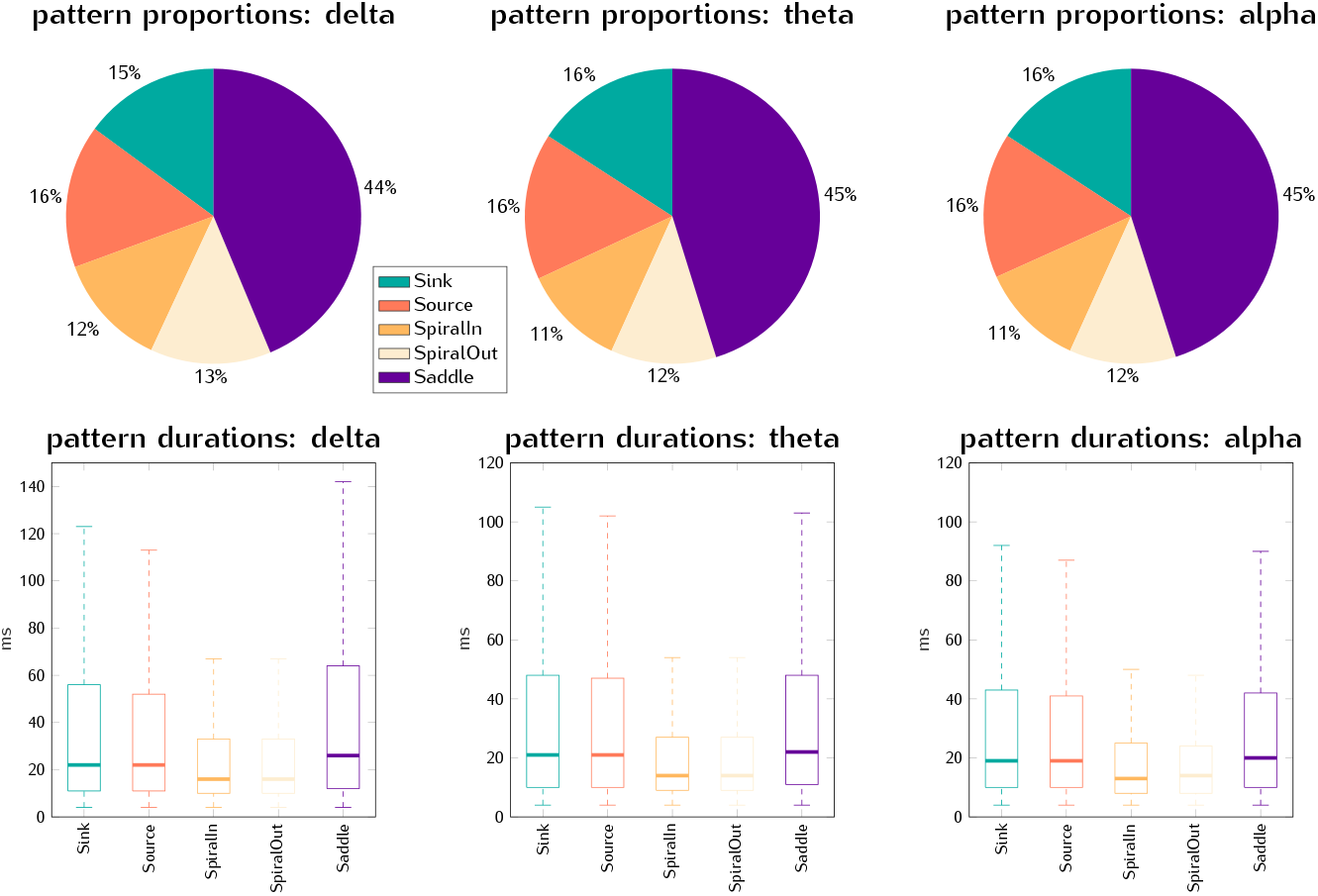
Proportion and duration of the five patterns identified in all the 16 subjects’ EEG: sink/source, spiral-in/spiral-out and saddle.

The ratios between the numbers of patterns varied very little across the three main frequency bands: *delta* [1, 4] Hz, *theta* [4, 8] Hz, or *alpha* [8,13] Hz.

We also evaluated only micro patterns covering a 2 × 2 grid patch (the whole projected 2D scalp grid is 67 × 67). For each event trial, which lasted 5000 ms, one can identify more than 10 patterns per millisecond, attesting to the highly heterogeneous EEG signal and complex brain/head geometry. These patterns last 20 to 40ms on average. Saddle patterns tend to last slightly longer than the others. However, in the very large number of patterns, certain patterns can last up to 1–2 seconds.

#### Location of patterns

With such a large amount of patterns and due to the complex head geometry, it is not possible to directly apply the same simple approach as for simulated data. We will not study every single pattern but focus on the brain regions where EEG activity is predominant and generated clusters of patterns.

A density (count per pixel) map can be obtained for each pattern in a trial. This map is Z-score normalized and only regions of interest with a pattern density at least two times the standard deviations above the mean overall density are kept. These pixels are traced and enclosed with isolines, forming the cluster regions. We consider the degree of overlap between the cluster regions of different patterns to be proportional to the spatial correlation between the patterns.

Figure 7 shows a representative example of this pattern cluster based approach (similar plots are obtained for other subjects). We took the mean area of the three given patterns as reference to calculate the overlap percentage. Clusters from sources and sinks overlap greatly. Spiral clusters are also correlated with sources/sinks, with overlaps ranging between 60 — 70%. This suggested that locations of source/sink/spiral correlate. However, saddle clusters have less than 1% overlap with source/sink. From the cluster density maps, we can see that saddle clusters are located between source/sink clusters. This way allows us to validate the hypothesis in real EEG data.

**Figure 7.**
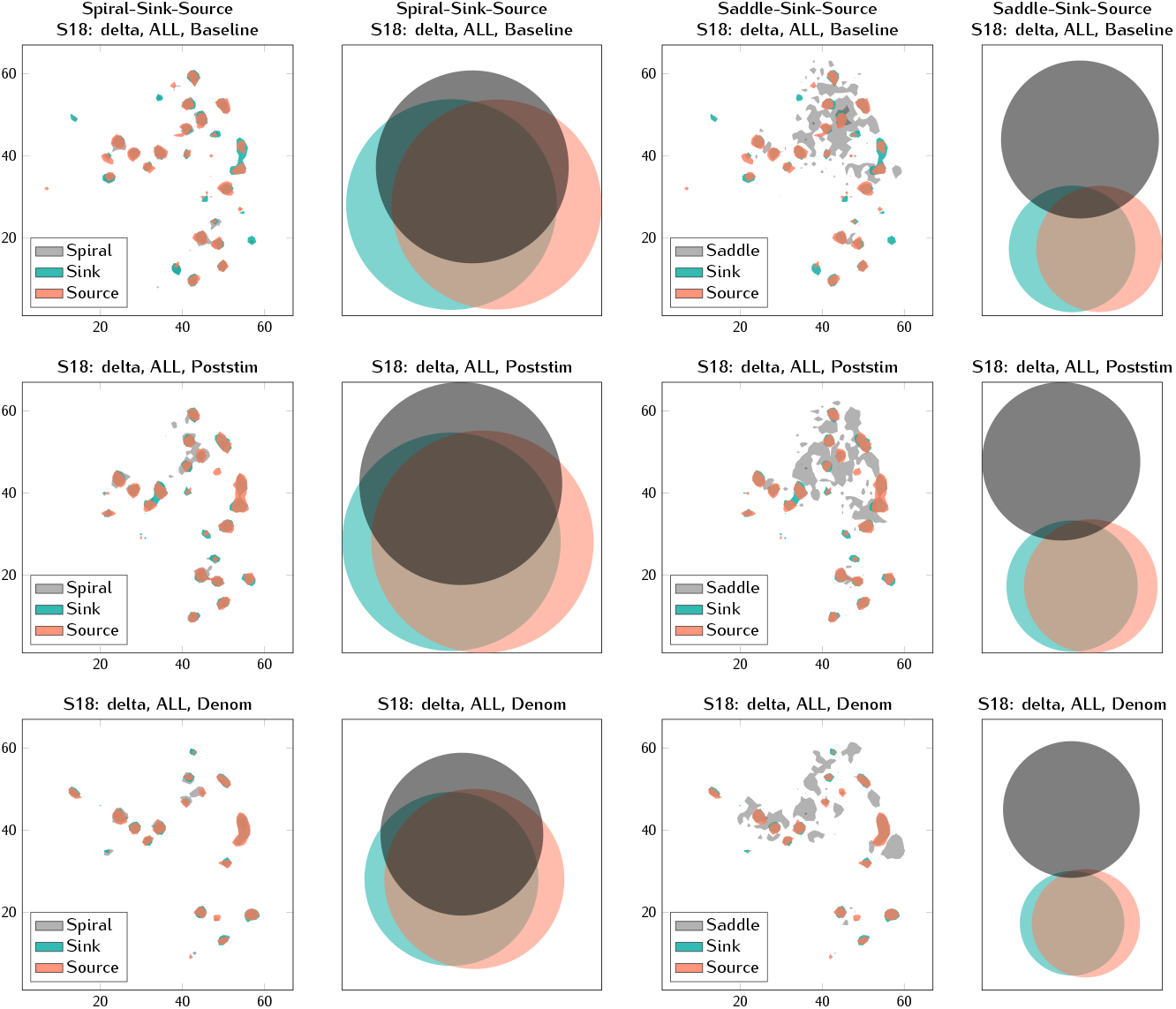
Two examples of pattern clusters in the real EEG data from Subject 18 (S18) performing the picture-naming task: 1) spiral-sink-source (two leftmost columns) and 2) saddle-sink-source (right two columns). In *each pair* of columns, the plots on the left are spatial plots of main thresholded pattern clusters across the entire scalp on a 67× 67 grid, and the column to the right counts (and overlapping) of clusters of the given patterns. Baseline ([-2s, 0s]) – period before onset of picture presentation; Poststim ([0s, 1.5s]) – after picture was presented; Denom ([1.5s, 3s]) – naming period. *ALL* means that patterns from all validated trials are combined.

Based on this approach, we are able to quantify the amount of overlap among the pattern clusters for each subject’s EEG data. Source and sink clusters generally overlap by 80 — 90%. We consider here only core intersections of three patterns (source-sink-spiral, and source-sink-saddle):

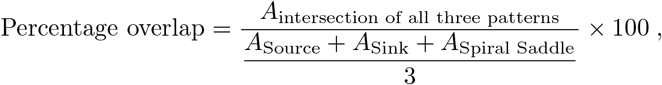

where *A* denotes area. Overall, as shown in Figure 8, source–sink–spiral cluster regions overlap with each other greatly, with median percentages around 60%, and hence have high spatial correlation. Clusters of source or sink overlap less than 1% with saddle clusters. This minimal overlap is coherent with the property of saddles that they are located between sources/sinks, and is also visually apparent in Figure 7, where most saddle clusters can be seen surrounded by source and sink clusters. Similar properties are observed for other subjects.

**Figure 8.**
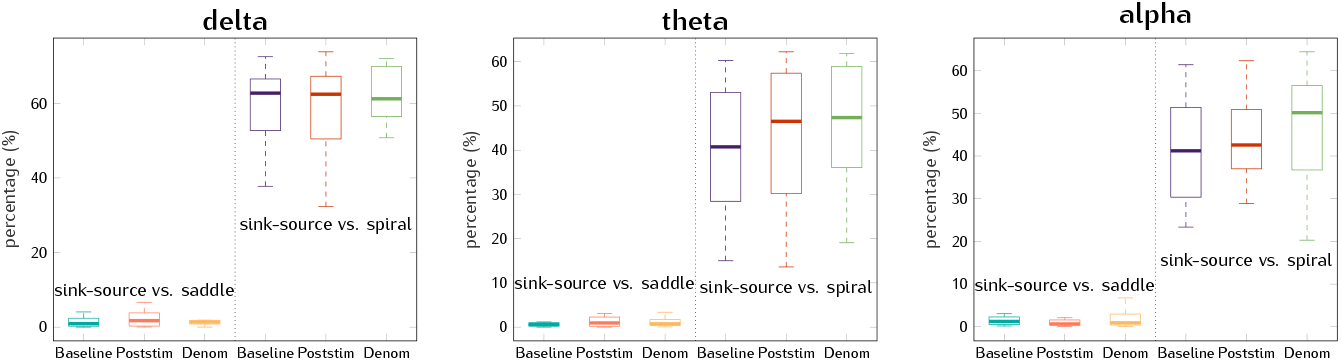
Percentage overlap by pattern clusters in three frequency bands for all subjects.

Location of patterns very weakly depends on frequency band. These observations hold for the three frequency bands except for the delta band, for which the overlap among source/sink/spiral is slightly higher than the other bands.

#### Number of patterns

Here we test the validity of Property 2, which states that the difference between the total number of all patterns except for saddles and the number of saddles is constant over time. For a finer-grained analysis, we plot the average frequency of patterns across epochs and over time for saddles alone as well as for the sum of all the other patterns, by subject, electrode, and frequency. Figure 9 shows the difference between the number of saddles and the sum of all other patterns during the picture-naming task considered for all subjects, trials, brain areas and frequencies. As stated in Property 2, this difference should be constant (positive, close to 0). The number of individual patterns (e.g., saddles) during naming is considered in the next section.

**Figure 9.**
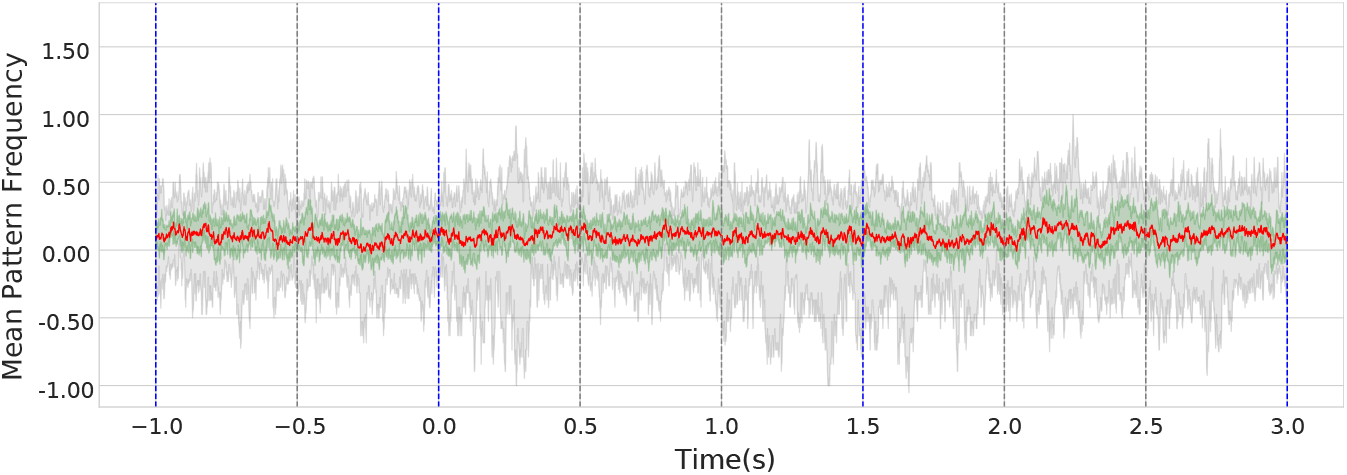
Mean pattern frequency difference between saddles and the sum of all other patterns across all subjects and all epochs in the real EEG data. The median of the pattern frequency (red curve) is close to constant, corroborating the hypothesis of constant difference (Eq. 1). The shaded green area delimits the first (bottom) and third (top) quartiles. The shaded gray area demarcates minimum and maximum frequencies.

## 4 Patterns during picture naming

The number of each type of patterns (i.e. source, sink, saddle, spiralin, spiral-out) was analyzed for each of the 16 subjects by EEG signal frequency band (delta, theta, alpha, beta) and by cortical zone (frontal, parietal, occipital, left temporal, right temporal). The 270 trials of picture naming were separated into two groups: a group of 70 trials with the single control word (*chien,* i.e. “dog”), and another group with the remaining 200 trials with all other words. Trials with signal artifacts were removed, leaving variable number of trials per subject. The average number of patterns for each group was counted in a time window starting 1 s (t0-1) before picture onset (time 0, t0) to the start of the naming prompt at t0+1.5 s. Signal amplitude and phase were analyzed independently.

The most regular frequency pattern during picture naming with the least variability was for “saddle” patterns in the delta frequency range for the word group “not-dog” (other words).

A typical example of the temporal evolution of the number of patterns is shown in Figure 10. The dependence of the number of patterns on time is similar for frontal and occipital zones and for other types of patterns (not shown). The number of patterns decreases at the beginning of epochs. In the box plots (Figure 10, right), the pattern frequency tends to decrease for the first 3 or 4 intervals in all subjects, with the minimum frequency reached by interval 3 or 4 for much subjects. For subjects where this was not the case, the frequency levels off or reduces in slope around the third and fourth intervals.

**Figure 10.**
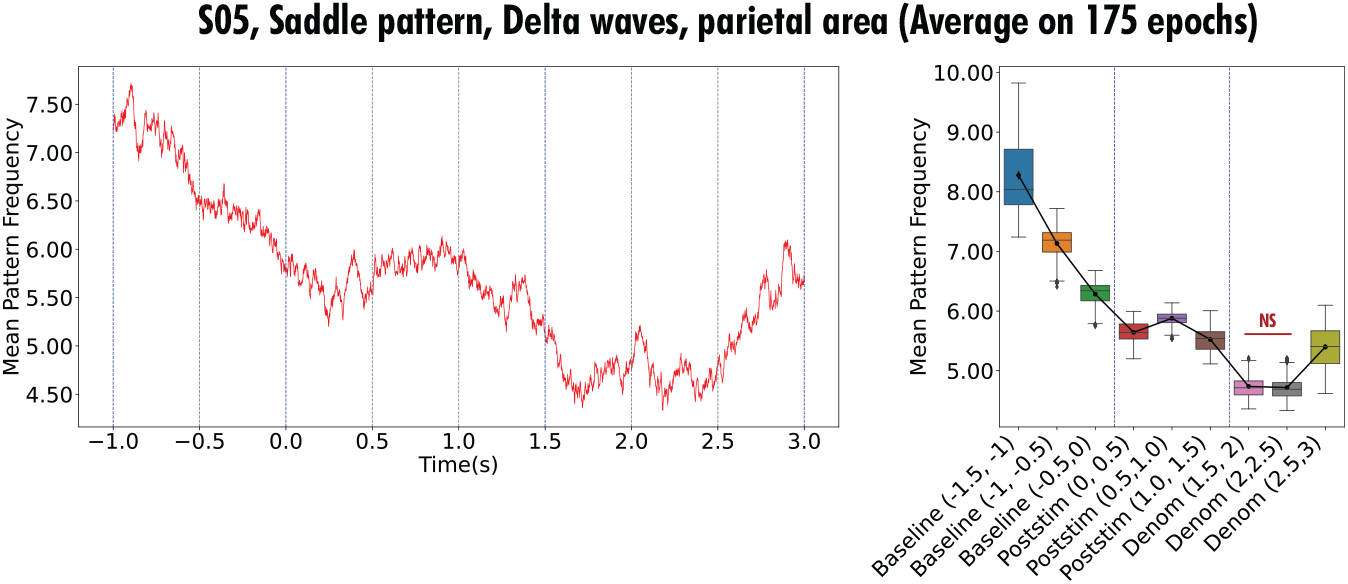
Temporal evolution of the number of saddle patterns from Subject 5 (S05) for the frequency delta range and non-control words (“not-dog”) in the parietal zone. The graph represents an average number of patterns with respect to all trials of this group. Red curve (left plot) shows the number of patterns averaged across 175 epochs in 5-ms time-bins. In the box plot (right), the black curve interpolates between the average number of patterns at each 500-ms interval.

This decrease of the number of patterns in the beginning of an epoch can be quantified for all subjects altogether by the following method. Let *N_p_*(*t_i_*) be the number of saddles at time *t_i_* in the parietal zone (Figure 10, panel C, red curve). Consider an average number of patterns with respect to three neighboring time points:

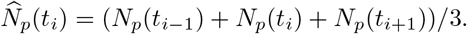

For the Heaviside function *H*(*x*), defined by the conditions *H*(*x*) = 1 for *x* > 0 and *H*(*x*) = 0 for *x* ≤ 0, we have

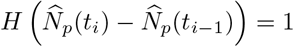

if 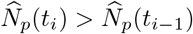, that is, if the function 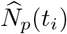 is increasing at this time point. Then the sum

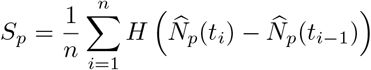

gives the proportion of time points where the function 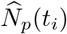 is increasing. Figure 11 presents violin plots showing the distribution across subjects of the percentage of increases for three event intervals: before picture presentation/visual stimulus (baseline), during visual stimulus but before naming, and during naming. The relatively small values for the first histogram correspond to the decrease in the number of patterns.

**Figure 11.**
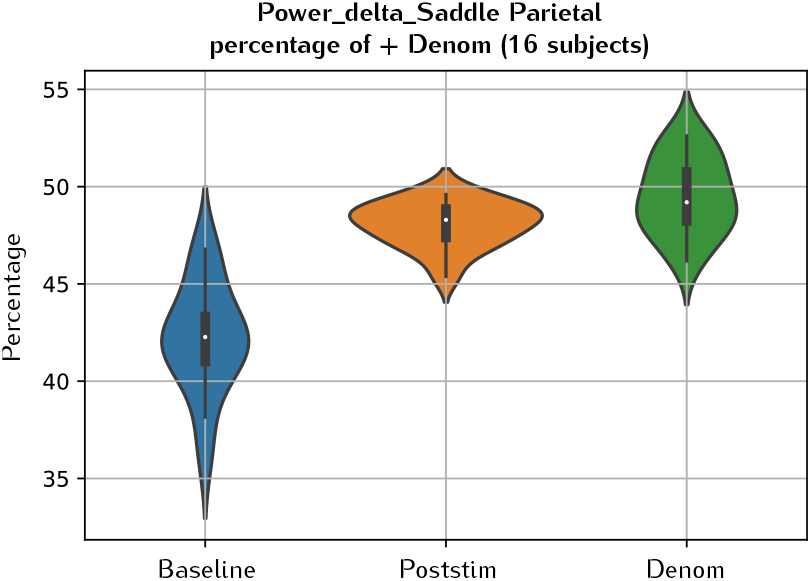
Distribution of the percentage increase of points across all subjects for three event intervals: before picture onset/visual stimulus (baseline), during visual stimulus but before naming (poststim), and during naming (denom).

The second event interval (before naming) is characterized by a weak increase (or plateau) of the number of patterns which further decreases, likely due to anticipating vocal articulation. This behavior can be observed in the individual curves for all subjects (e.g., Figure 10) but not for group averages (Figure 11) due to the weakness of the effect.

The number of patterns as a function of time across an epoch depends on frequency range. Typical examples of such dependence for the alpha and beta frequency bands are shown in Figure 12 for the same subject. The number of patterns has a tendency to decrease towards halfway through the epoch in the alpha band but a tendency to increase in the beta band. This behavior is somewhat generalizable to other subjects. We will discuss possible interpretations of these results in the next section. The results for other frequency ranges and for phase patterns are not presented here for brevity. All results presented in this section concern the group of words different from the control word “dog”. The results are similar for the other group of words containing the repetition of the control word.

**Figure 12.**
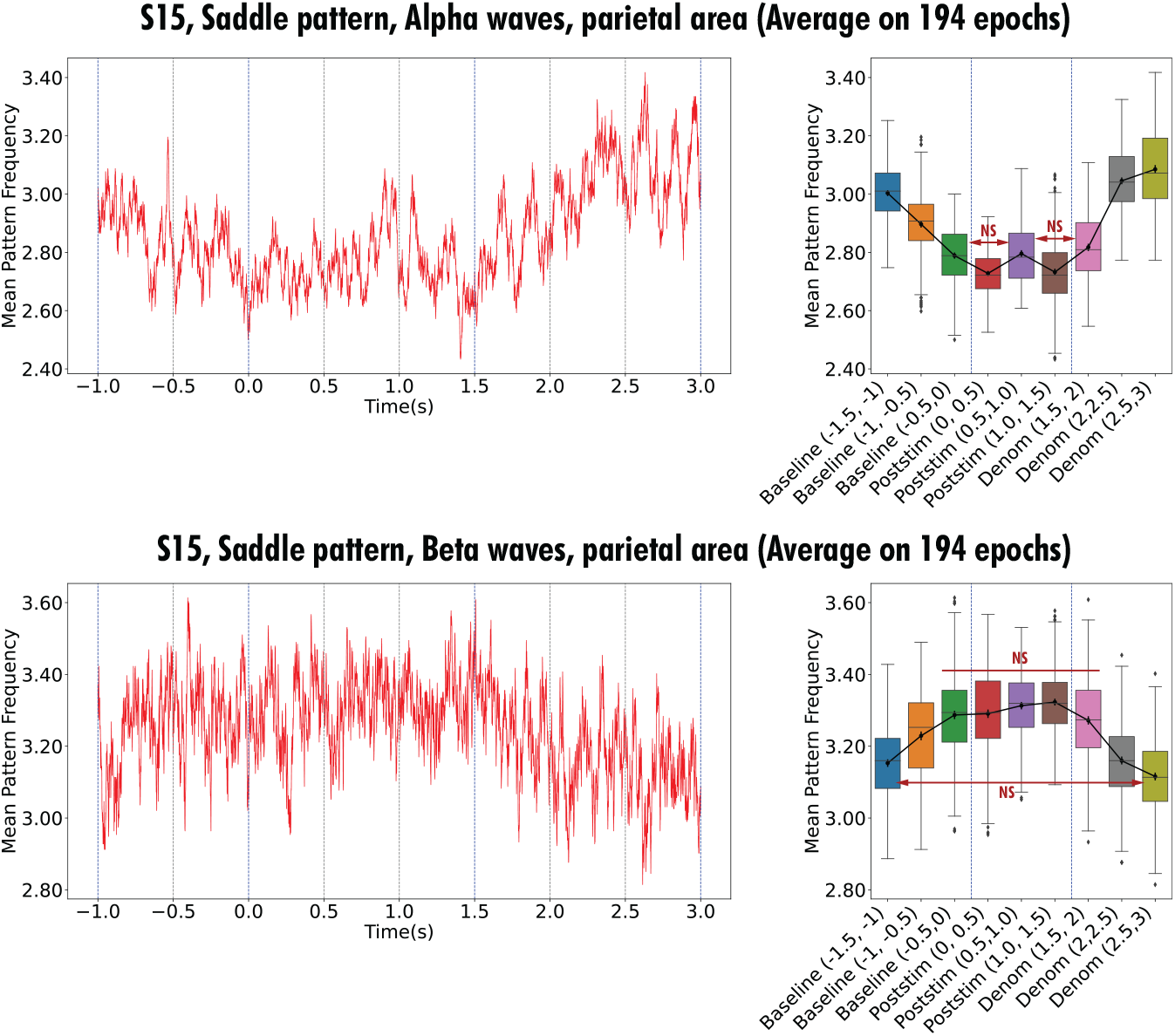
Temporal evolution of the number of saddle patterns in one subject (S15) in the frequency alpha / beta band for the non-control word group (“not-dog”) in the parietal zone. The graph represents an average number of patterns with respect to all trials of this word group. Red curve (left plot) shows the number of patterns averaged across 175 epochs in 5-ms time-bins. In the box plot (right), the black curve interpolates between the average number of patterns at each 500-ms interval.

## 5 Discussion

### 5.1 Spatiotemporal dynamics

#### Brain micro-states and sources

One of the main advantages of EEG data is its very high temporal resolution, which makes it possible to capture fast brain dynamics. The spatiotemporal dynamics in EEG data can be characterized by time-dependent amplitude and phase changes. These dynamics have been found useful in describing different brain states (rest, cognitive tasks, motion) [19,20] or disorders (e.g., aphasia, epilepsy, schizophrenia) [17,24].

Brain micro-states are often defined as relatively stable (weakly changing) distributions of electric potential observed during sufficiently long time intervals (tens of millisecond) with rapid transitions between them [17]. Several dominant micro-states can cover an essential part of dynamics during observation time window. To simplify the analysis of micro-states, they can be commonly characterized by the maximum and minimum of the potential distribution (e.g., direction of the interval connecting them) [16,34] or by the time trajectory of its maximum [27].

Time sequences of micro-states give a rather complete representation of spatiotemporal dynamics, though they do not seem to capture some dynamic effects, such as travelling waves (plane, rotating), or some specific types of dynamics (sources, sinks, saddles). Properties of brain micro-states are related to the underlying brain sources. From the biological point of view, brain sources are determined by cation flux from the intracellular space to the extracellular space, and brain sinks to the inverse flux [35]. Assumption that the brain is electrically neutral implies that sources and sinks have the same intensity. In simplified models, they are considered pairwise and close to each other (dipole). However, positive and negative poles of the dipole can be distant [35]. The distribution of electric potential at the surface is determined by the dipole position and orientation. The maximum and the minimum of the potential distribution do not necessarily correspond to the dipole location.

Identification of brain sources for each micro-state and their comparison with fMRI images allow the determination of the corresponding anatomic structures and to associate micro-states to brain functions [18–20]. The inverse problem of source identification has multiple solutions. The choice between them is to some extent arbitrary and can be determined by some additional factors (fMRI, anatomical structures). Thus, spatiotemporal dynamics of EEG data characterized by brain micro-states are determined by the change of brain sources, but the underlying regimes (patterns) are not yet identified. We consider in this work some of these regimes determined by phases and frequencies of brain sources.

#### Regimes determined by phases and frequencies of brain sources

Since spatiotemporal dynamics in EEG data is complex and depends on many factors, one needs to first identify some basic spatiotemporal regimes. We have shown here that the phase and frequency of the signal, for a given EEG signal frequency band (e.g. alpha) give rise to certain regimes in the spatiotemporal dynamics induced by brain sources and their interactions. For the case of single positive and negative poles (source and sink), only standing waves are possible. We have observed this regime for different frequency bands, but it is more difficult to identify in a broad frequency band such as 2–40 Hz, due to the superposition of different frequency components in the signal.

The assumption of the electric neutrality of the brain implies that phase and frequency of the sources are the same if there are only two sources. In the case of three sources, this is not necessarily the case. We have three additional basic spatiotemporal regimes on top of standing waves.

If frequency and phase are the same for all three sources, like with two sources, the corresponding regime is a standing wave. However, if there are different phases and/or frequencies, the corresponding dynamics will be represented by a combination of standing and moving waves. These regimes are related to tACS modelling [36], and we have shown that simulation of electrical potentials under tACS stimulation (Supplementary Materials, B) also have dynamics that follow these regimes.

In the case of three sources with different phases and the same frequency (with constraints imposed but electric neutrality), we observe out-of-phase waves with periodically changing phases. If the frequencies are different, these are modulated waves with a periodically changing amplitude.

#### Micro-states and waves in basic regimes

These basic regimes also have characteristic micro-states. Standing waves have two micro-states (Figure 2 in Section 2) with periodic transitions between them (and time-dependent amplitude). In our simulated out-of-phase waves, there are three microstates (not shown). Each micro-state slowly varies with the position of the maximum of the potential distribution gradually changing. These maxima jump to distance locations during transitions between micro-states.

Similar micro-states are observed for generated data in the case of different frequencies. However, one important difference is that the rotation changes directions every half a period.

There are two types of moving waves in basic regimes. The first one is determined by slow variations in basic states. It can be observed as a motion of the maximum of the potential distribution (trajectory of the yellow dots in Figure 3).

The second type of moving waves is related to the transition between micro-states since it is not instantaneous. The combination of both wave types can produce the rotating waves as can be seen in Figure 3, from simulated data. We have observed similar regimes in resting-state EEG data from human subjects (not shown) (see [20]).

It is important to indicate that in the model considered here, oscillations in the EEG data are measurements of internal brain sources on the outer surface (cortex) and not direct measurements of the neuronal electric activity in the cortex. As indirect justification of this hypothesis, we note that these waves (speed, direction) are not apparently influenced by sulci and gyri, which would be the case if they propagate along the cortex. Moreover, their speed is of the order of meters per second, while the speed of electric impulses in unmyelinated axons in the cortex grey matter is about ten times less. Therefore, if these waves appear only as projection of brain sources, they may not function for the synchronization of distant brain areas but indicate synchronization of distant brain sources.

### 5.2 Optical flow patterns

#### Location and number of patterns

The three regimes discussed above (standing waves, out-of-phase waves, and modulated waves) are observed in the analytical solution (Supplementary Materials, C), in simulated data, and in real EEG data. Analyzing the properties of optical flow patterns for the theoretical solution, we can expect that similar properties hold for both generated data and real EEG data, since spatiotemporal regimes for them are similar.

Analysis of optical flow patterns for standing waves shows that sources and sinks alternate in occupying the same locations and that this location does not depend on time. Moreover, saddles are located between sinks and sources, with their frequency approximately equal to the total frequency of sources and sinks. All these properties are confirmed for the simulated and EEG data, as well as for all three regimes.

Another conclusion from the theoretical analysis is that complex patterns and travelling plane waves are mutually exclusive. This result corroborates with previous reports [29,30]. Verification of this in EEG data is beyond the scope of this work.

#### Spatiotemporal patterns and word naming

We have determined some correlations in the picture-naming task and the number of observable spatiotemporal patterns. The most stable behavior across subjects was observed in the delta range; in this range, the number of patterns decreases in the beginning of the epoch. Such a decrease can also be observed in the alpha range, but inter-subject variation is greater. In contrast, the number of patterns in the beta range has the tendency to increase towards the middle of epoch.

The number of patterns decreases in the delta range, and this decrease begins before picture presentation (Figure 10). Therefore, we can conjecture that the number of patterns might indicate the effect of anticipation known for delta rhythms (see [37], page 52 and the references therein). If this anticipation down-regulates activity of some brain sources, then it can manifest as a decrease in the number of patterns. Similarly, the second (smaller) decrease is observed at vocalization onset during naming, possibly during anticipating word pronunciation. Alpha rhythms have been implicated in inhibition ([37], pages 46-47) and could be acting on brain sources, leading to the decrease of the number of patterns (Figures in 12). Furthermore, it is known that alpha and gamma rhythms can be complementary ([37], page 47). We observe a possible complementary interaction between alpha and beta rhythms (Figures in 12). Finally, there is a possible correlation between theta-rhythm amplitude and delta-rhythm phase with the number of patterns for some subjects (not shown).

Let us also note that picture recognition is accompanied by a larger ERP amplitude. However, as suggested from our theoretical analysis here, the amplitude of oscillations alone does not influence the number of optical flow patterns. Therefore, activation and/or inhibition between brain sources during cognitive activity is very likely a driving force in spatiotemporal dynamics, which determines these patterns. Changes to the phase and frequency of oscillations from different sources can influence the number and dynamics of patterns, as is seen for the three main regimes, and can arguably be a means to influence communication between sources with possible observable functional behavioral changes.

## Acknowledgments

The authors thank Professor Monica Baciu and Sylvain Harquel from LPNC at the University of Grenoble Alpes (UGA) for their collaboration and help with data collection.

## Appendix A EEG data acquisition and treatment

### A.1 Data collection

Sixteen native French-speaking men aged 18—70 years participated in the Picture Naming Task study. Inclusion criteria were normal (or corrected to normal) vision and hearing, and right-handedness as assessed by a handedness questionnaire [38]. Exclusion criteria were any history of neurological or psychiatric disorders, drug addiction, or head trauma. Pictures for the task were taken from the Snodgrass & Vanderwart black-and-white line drawing corpus [39]. Pictures were shown on a screen. The subject’s voice was recorded and synchronized with EEG (96 EEG channels with sampling frequency 1 kHz). The study was approved by the Research Ethics Committee CER Grenoble Alpes (Avis-2020-09-01-3).

### A.2 Data preprocessing and analysis

The raw signal from the 96 channels for all 270 trials were first epoched with duration 5.5 seconds (2 seconds pre- and 3.5 seconds post-visual stimulation onset), and baseline corrected ([-1s, 0s]). Bad epochs were removed (e.g. eye blinks, eye or head movements), and the remaining epochs were band-pass filtered at 0.1—40Hz.

The preprocessed EEG signals with three-dimensional sensor coordinates were then projected onto a two-dimensional scalp plane for selected time points. Values between electrodes were interpolated using biharmonic splines [40], resulting in a 67 × 67 grid. Topographical scalp maps were created in this way from the signal from the 96 channels for all 5500 samples.

### A.3 Pattern extraction pipeline

To identify the (2D) pattern types: *saddle, spiral-in, spiral-out, sink* and *source,* critical points in the vector fields derived from the EEG signals were identified. Vector fields were obtained by computing the optical flow from the analytical phase or amplitude, which were extracted from the pre-processed EEG signals using the Hilbert transform (planar projection in a grid).

The detected patterns are then analyzed via different techniques taking in consideration multiple characteristics such as frequency, spatial area, and observation period.

The pattern extraction pipeline can be summarized as follows:

1. **Signal time-frequency analysis**: extract phase and power of signals using the Hilbert transform. The Hilbert transform [41] extracts the instantaneous phase and amplitude from these four frequency bands of the preprocessed EEG data: `delta` (1—4Hz), `theta` (4—8Hz), `alpha` (8—13Hz), and `beta` (13—30Hz).
2. **2D projection**: 2D projection of the signal on the scalp for each time frame. The spatial distribution of the power of the EEG signal is visualized by planar projection of electrode position coordinates on the scalp. These topographic maps were smoothed with *biharmonic spline interpolation* [40]. ^2^.
3. **Optical-flow analysis**: identification of the dynamics of the signal by calculation of vector fields (between two time points) with the Horn-Schunck method [43].
4. **Pattern identification**: Identify patterns in the vector fields (e.g. sinks, sources). NeuroPatt [29] was used to identify patterns on the preprocessed EEG data.

This resulted in 270 × 5500 = 1.485M projections (topographic maps).

## Appendix B EEG data simulation with SimNIBS

SimNIBS [44] was used to generate synthetic EEG data using a realistic head model. Three different 3-electrode tDCS stimulations were simulated, and the generated signals combined with time-dependent weights in order to simulate tACS [32].

### B.1 Simulation of tDCS

The example head model [45] and the default electrode positions were used. The three setups differed by input currents, as listed in Table S1. The location of the stimulation electrodes were the same for all simulations (AFz, POz, C5).

**Table S1.**
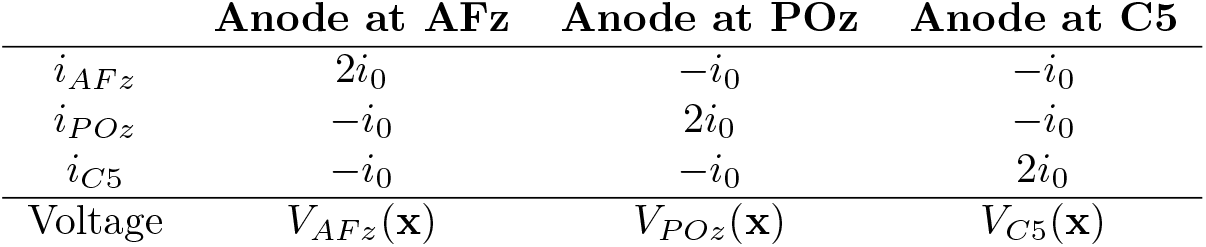
The input current at the three stimulation electrodes for the three tDCS stimulations. *i*_0_ = 500 *μA*.

|  | Anode at AFz | Anode at POz | Anode at C5 |
| --- | --- | --- | --- |
| $i_{AFz}$ | $2i_0$ | $-i_0$ | $-i_0$ |
| $i_{POz}$ | $-i_0$ | $2i_0$ | $-i_0$ |
| $i_{C5}$ | $-i_0$ | $-i_0$ | $2i_0$ |
| Voltage | $V_{AFz}(\mathbf{x})$ | $V_{POz}(\mathbf{x})$ | $V_{C5}(\mathbf{x})$ |

The current *i*_0_ = 500 *μA* is a quarter of the maximum stimulation intensity considered safe [46]. Each simulation results in an exogenous electric field where the voltage at each point **x**, *V_elec_*(**x**) is computed at approximately 250000 points **x** of the SimNBS head-mesh grid, identified by their 3D coordinates inside the “brain”. Figure S1 shows the results of the tDCS simulation with anode over POz.

### B.2 Simulation of tACS

The results of simulated tDCS stimulations were then combined into simulations of tACS. Additional care has to be made because the stimulations are out of phase.

**Figure S1.**
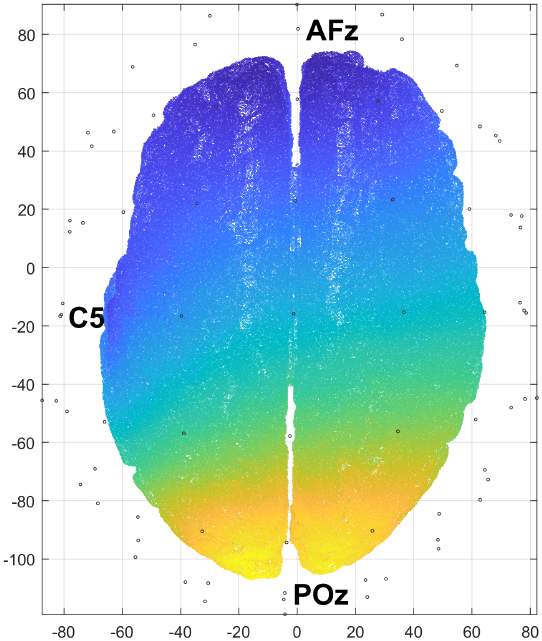
One simulation of tDCS with positions of electrodes in 3D.

The input stimulation currents were denoted by:

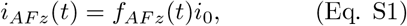

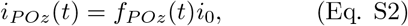

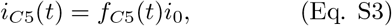

such that

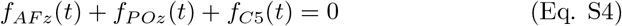

for all times *t*. Kirchhoff’s Current Law (Eq. S4) is a necessary condition for physical or simulated stimulation [33], and is analogous to (Eq. S9).

The tDCS simulations were assigned normalized weights (*α*(*t*), *β*(*t*), *γ*(*t*)), such that the point 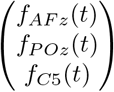 is the barycentre of the points

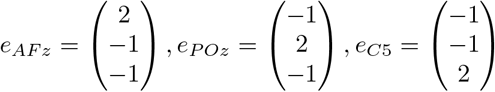

with these weights. This condition is equivalent to the system:

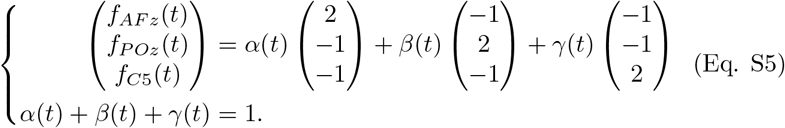

Note that the system (Eq. S5) has a unique solution since all four points belong to the same 2D plane (Eq. S4) of admissible current intensities, and the three points *e_AFz_, e_POz_, e_C5_* are not on the same affine line.

The system (Eq. S5) can be solved using linear algebra. The formulae for the weights (*α*(*t*),*β*(*t*),*γ*(*t*)) are the following:

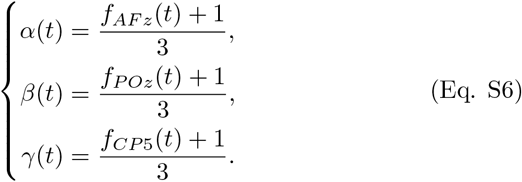

One can check that these values satisfy both equations of the system (Eq. S5) provided that the input functions satisfy (Eq. S4).

The solution of (Eq. S5) can be applied to the results of three tDCS simulations in order to simulate tACS. Indeed, the tDCS was simulated with the input currents equal *e_AF_z__ i*_0_, *e_PO_z__ i*_0_ and *e*_*C*5_*z*__*i*_0_. By (Eq. S5) and the assumption of linearity, the (exogenous) voltage generated by the tACS stimulation equals:

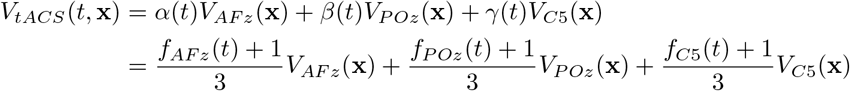

The above procedure allows computing the simulated voltages for tACS simulation for any input currents determined by the functions *f_elec_*(*t*). It was applied to simulate the following types of tACS (see Subsection 2.1).

a. the currents at *AFz* an *POz* are equal, and their frequency equals *f*_1_ = 9*Hz*;
b. the phase of the electrode *POz* is posterior to AFz by *ϕ*_2_ = 138°;
c. the frequency of *POz* equals *f*_2_ = 10*Hz*.

Table S2 contains the formulas being used. The intensity at *C*5 (return electrode) was adjusted in order to satisfy (Eq. S4).

**Table S2.**
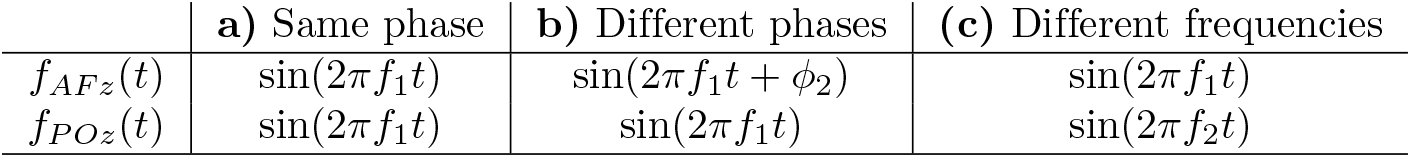
The three tACS stimulations.

|  | a) Same phase | b) Different phases | c) Different frequencies |
| --- | --- | --- | --- |
| $f_{AFz}(t)$ | $\sin(2\pi f_1 t)$ | $\sin(2\pi f_1 t + \phi_2)$ | $\sin(2\pi f_1 t)$ |
| $f_{POz}(t)$ | $\sin(2\pi f_1 t)$ | $\sin(2\pi f_1 t)$ | $\sin(2\pi f_2 t)$ |

### B.3 Restriction to the EEG electrodes

The algorithm above always allows computing the voltages at all points of the grid. Extracting the voltages at a small number of electrodes on the scalp surface is necessary in order to create a dataset with parameters comparable to experimental EEG data.

Coordinates of 30 electrodes were chosen from the list of 76 items provided with the example dataset [45]. Their coordinates (3D) were used for estimating the tACS voltages (by using the voltage at the closest point of the grid as an estimation of the voltage at an electrode).

## Appendix C Analytical solution

### C.1 Model of brain sources and approximate solution

Consider a 3D domain Ω representing human brain with its surface corresponding to the brain cortex. The distribution of electric potential in this domain can be described by the Poisson equation:

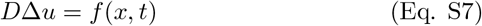

where the source *f* (*x, t*) depends on time. Therefore, solution *u*(*x,t*) of this equation also depends on time as parameter. If we consider no-flux boundary condition at the boundary,

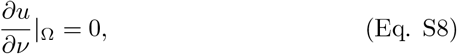

where *ν* is the outer normal derivative vector, then problem (Eq. S7)–(Eq. S8) has a solution if and only if

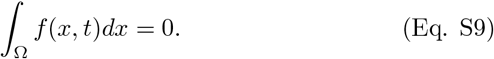

It is a classical solvability condition of elliptic boundary value problems, and it is conventionally used in neuroscience in electric brain stimulation [36].

Consider point-wise sources represented by *δ*-function with time-periodic amplitude taken, for certainty, as sin(*kt*). In the case of two sources located at points 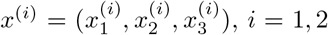, condition (Eq. S9) implies that they have opposite phase:

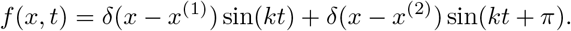

Then solution *u*(*x,t*) of problem (Eq. S7)–(Eq. S9) can be written as follows:

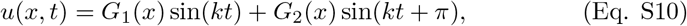

where *G_i_*(*x*), *i* = 1, 2 are the corresponding Green’s functions. If the sources are sufficiently far from the boundary of the domain, then the functions *G_i_*(*x*) can be approximated by Green’s function in the whole space:

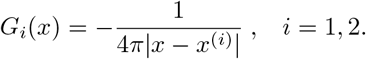

This solution can be easily generalized for any number of sources.

**Figure S2.**
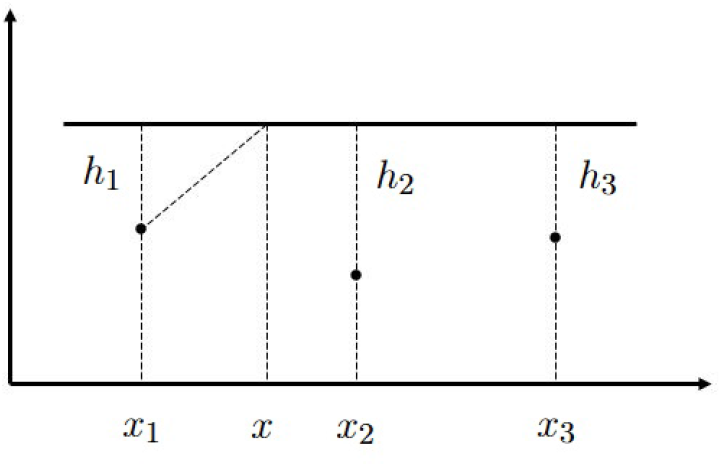
Cross-section of the domain Ω with three EEG sources and the boundary of the domain a straight line above them.

We will study dynamics of solutions of problem (Eq. S7), (Eq. S8) in the case of three point-wise sources:

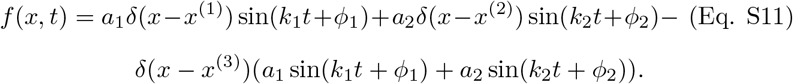

Let us note that function *f* (*x,t*) is written here in such form that condition (Eq. S9) is satisfied for any values of *a*_1_, *a*_2_, *k*_1_, *k*_2_ and source locations *x*^(*i*)^, *i* = 1,2, 3. Replacing Green’s functions in the bounded domain by the corresponding Green’s functions in the whole space, we approximate solution of problem (Eq. S7), (Eq. S8) by the following function:

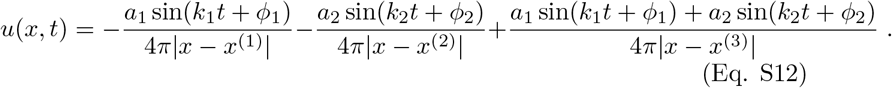

In order to more easily assess the properties of this solution, consider the plane passing through the points *x*^(*i*)^, *i* = 1, 2, 3 and suppose, for simplicity of presentation, that the intersection of this plane with the boundary of the domain Ω is a straight line (Figure S2). Thus, we consider a cross-section of the 3D domain by a plane and introduce 2D coordinates (*ξ, η*) with the coordinates (*ξ_i_,η_i_*) of the sources and distances *h_i_* from the sources to the boundary. Then solution (Eq. S12) can be written as follows:

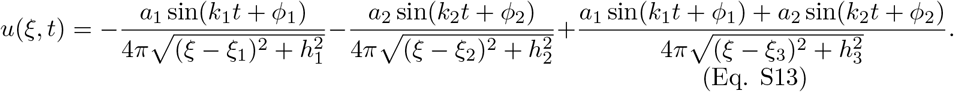

Here is the coordinate of the point at the boundary of the domain.

**Figure S3.**
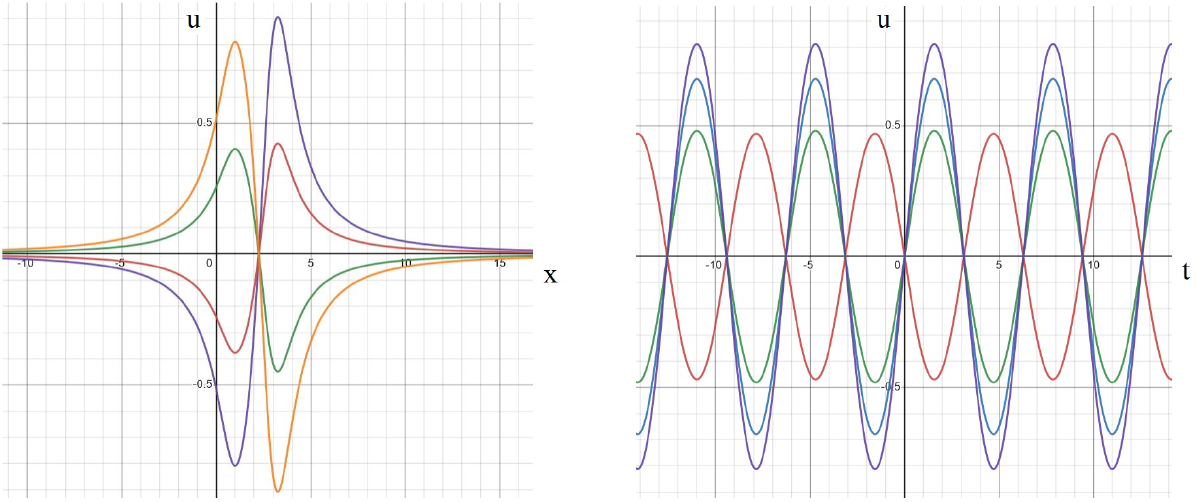
Solution (Eq. S13) in the case of equal frequencies and phases as a function of space variable for different moments of time (left). The same solution as a function of time at different space points (right). The values of parameters: *k*_1_ = *k*_2_ = 1, *a*_1_ = *a*_2_ = *a*_3_ = –4*π*, *h*_1_ = *h*_2_ = *h*_3_ = 1, *ξ*_1_ = 1, *ξ*_2_ = 2, *ξ*_3_ =3, *ϕ*_1_ = *ϕ*_2_ = 0. Left: consecutive moments of time: *t*_1_ = 0(*blue*), *t*_2_ = 2(*green*), *t*_3_ = 3(*orange*). Right: time dependence at different space points: *ξ* = 2.6(*red*), *ξ* = 1.8 (*green*), *ξ* = 1.5(*blue*), *ξ* = 1(*violet*).

### C.2 Standing waves and out-of-phase dynamics

We consider dynamics of solution (Eq. S13) in the following cases: equal frequencies and phases, equal frequencies and different phases, different frequencies and equal phases, different frequencies and phases. In the first case, for *k*_1_ = *k*_2_ and *ϕ*_1_ = *ϕ*_2_, this solution describes standing waves (Figure S3). Potential distribution in space for a fixed moment of time is positive in one half-axis and negative in the other one. They oscillate in time alternating positive and negative values, but the boundary between them where *u*(*x,t*) = 0 does not depend on time. Furthermore, the maximum and the minimums of the solution can have only two possible space locations periodically jumping between them. These fixed position of the maximum during half-period and jumps to another fixed position are specific for standing waves.

If we fix the space point and consider the solution as a function of time (Figure S3, right), we observe a periodic function with a constant amplitude. By analogy with EEG data, such functions can be interpreted as signals registered with different electrodes. Different space points correspond to different electrodes. There is precise synchronization of these signals, they all vanish at the same moments of time.

**Figure S4.**
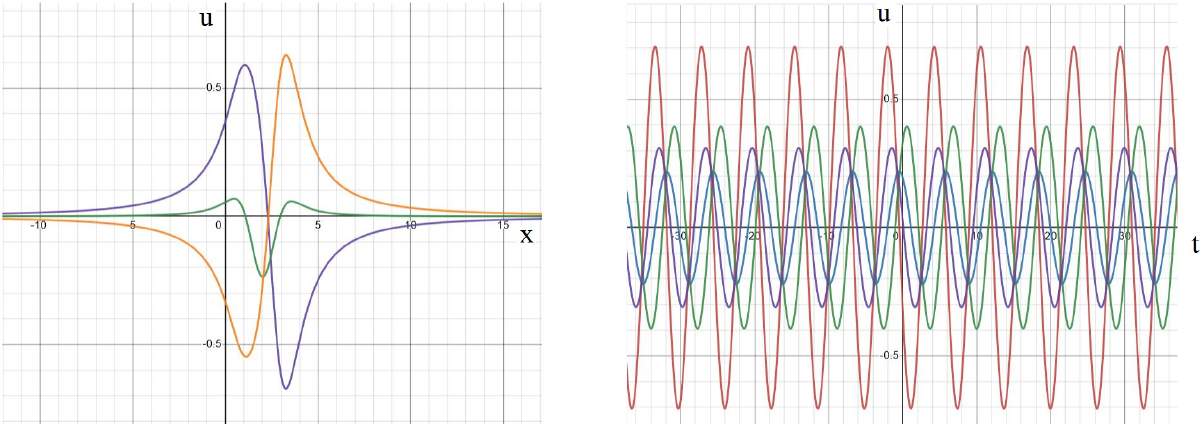
Solution (Eq. S13) in the case of equal frequencies and different phases as a function of space variable for different moments of time (left). The same solution as a function of time at different space points (right). The values of parameters: *k*_1_ = *k*_2_ = 1, *α*_1_ = *α*_2_ = *α*_3_ = –4*π*, *h*_1_ = *h*_2_ = *h*_3_ = 1, *ξ*_1_ = 1, *ξ*_2_ = 2, *ξ*_3_ = 3, *ϕ*_1_, = 0, *ϕ*_2_ = 1.3*π*/3. Left: consecutive moments of time: *t*_1_ = 0(*blue*), *t*_2_ = 2(*green*), *t*_3_ = 3(*orange*). Right: time dependence at different space points: *ξ* = 3.1(*red*), *ξ* = 2.2(*blue*), *ξ* = 1.8(*green*), *ξ* = 1(*violet*).

In the case of different phases and equal frequencies, the main dynamics of solution resemble standing wave, but the zero of solution is now time-dependent and it can be non-unique (Figure S4, left). Time dependence of solution at different space points are shifted in phase (Figure S4, right). The amplitudes of these signals remain constant in time. To fix the terminology, we call such solutions out-of-phase standing waves.

**Figure S5.**
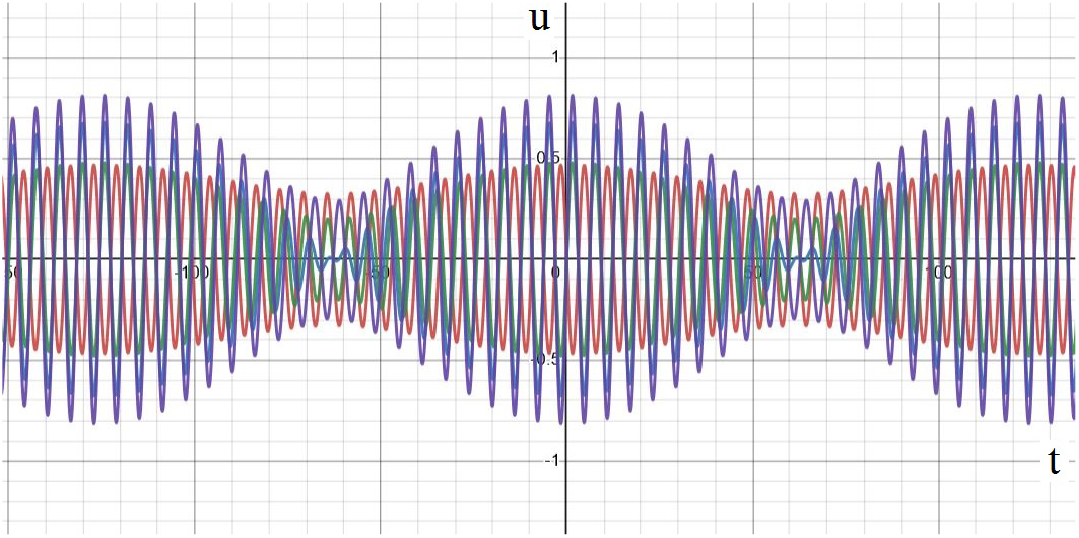
Solution (Eq. S13) in the case of different frequencies and equal phases as a function of time at different space points. The values of parameters: *k*_1_ = 1, *k*_2_ = 1.05, *a*_1_ = *a*_2_ = *a*_3_ = –4*π*, *h*_1_ = *h*_2_ = *h*_3_ = 1, *ξ*_1_ = 1, *ξ*_2_ = 2, *ξ*_3_ = 3, *ϕ*_1_ = *ϕ*_2_ = 0, time dependence at different space points: *ξ* = 3.1(*red*), *ξ* = 2.2(*blue*), *ξ* = 1.8(*green*), *ξ* = 1(*violet*).

If the phases are equal but the frequencies are different, we obtain modulated standing waves (Figure S5). The amplitude of signals changes periodically in time, while their phases are basically the same except for some transition zones. Such modulated signals arise due to addition of two periodic functions with different frequencies. For example, sin(*k*_1_*t*)+sin(*k*_2_*t*) gives a high frequency oscillation corresponding to (*k*_1_ + *k*_2_)/2 and low frequency modulation corresponding (*k*_1_ — *k*_2_)/2, assuming that *k*_1_, *k*_2_ > 0.

**Figure S6.**
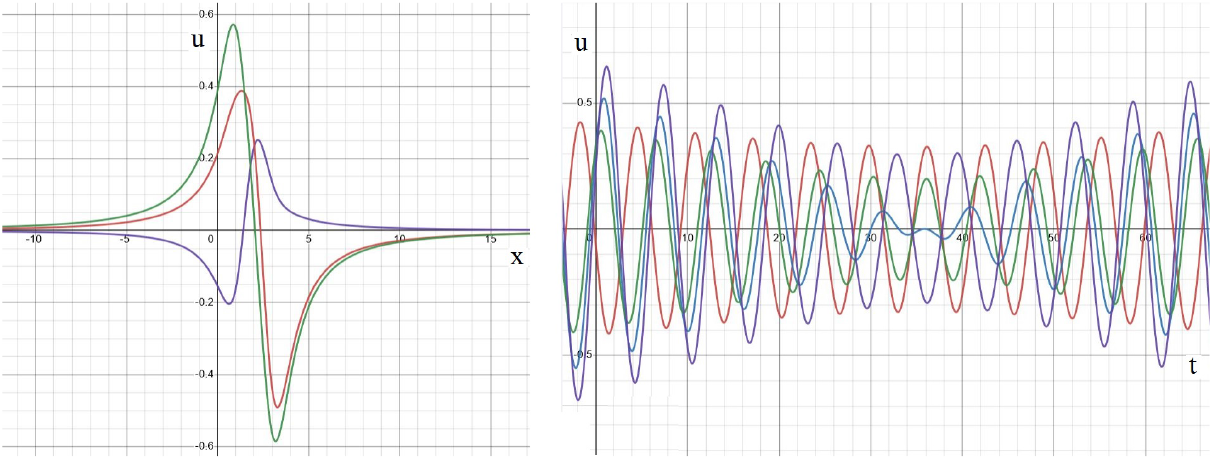
Solution (Eq. S13) in the case of different frequencies and phases as a function of space variable for different moments of time (left). The same solution as a function of time at different space points (right). The values of parameters: *k*_1_ = 1, *k*_2_ = 1.05, *a*_1_ = *a*_2_ = *a*_3_ = – 4*π*, *h*_1_ = *h*_2_ = *h*_3_ = 1, *ξ*_1_ = 1, *ξ*_2_ = 2, *ξ*_3_ = 3, *ϕ*_1_ = 0, *ϕ*_2_ = 1.3*π*/3. Left: consecutive moments of time: *t*_1_ = 0(*blue*), *t*_2_ = 2(*green*), *t*_3_ = 3(*orange*). Right: time dependence at different space points: *ξ* = 3.1(*red*), *ξ* = 2.2(*blue*), *ξ* = 1.8(*green*), *ξ* = 1(*violet*).

Let us finally consider the case of different frequencies and phases. In this case we obtain out-of-phase modulated waves (Figure S6) with time-dependent amplitude and shifted phase. Let us note that the maximum of solution sometimes moves in space as a function of time (Figure S6, left). From this point of view, we can characterize this solution as a combination of standing waves and travelling waves, though neither of them exactly corresponds to the strict definition of such waves.

## Appendix D Optical flow patterns for standing and travelling waves

The flow field in the optical flow method is described by the following system of equations:

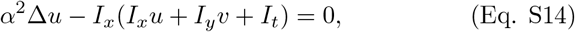

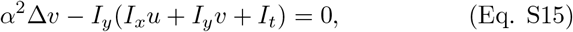

where *I*(*x,y,t*) is a given function which determines the flow field, the subscripts denote its partial derivatives, *α* is a regularization constant. We will present here some model examples illustrating the properties of the flow field depending on the function *I*.

### Linear function

Consider a linear function

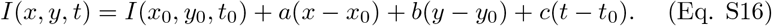

We look for the solution of system (Eq. S14), (Eq. S15) in the form *u* = *u*_0_, *ν* = *ku*_0_. If *b* ≠ 0, then *k* = – (*au*_0_ + *c*)/(*bu*_0_). A similar expression can be obtained if *a* ≠ 0. Hence, linear function *I* gives a constant vector field.

Strictly speaking, solving system (Eq. S14), (Eq. S15) in a bounded domain, we need to specify the boundary conditions. Since we are interested in local behavior of solution, we can choose boundary conditions in such a way that constructed solution satisfies them. If the function *I* is nonlinear, we can consider function (Eq. S16) as its linear approximation. The corresponding vector field is constant in the first approximation. Though it is not unique (up to a choice of *u*_0_ ≠ 0), it does not contain singular points.

An interesting particular example of function (Eq. S16) corresponds to linearization of travelling wave:

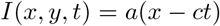

(propagating in the *x*-direction). In this case, *u* = *c, v* = 1, that is, horizontal component of the flow field equals the wave speed *c*. As before, there are no singular points of the flow field.

### Quadratic function

Consider the function

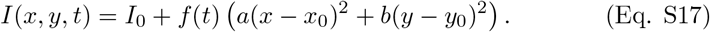

It can be considered as an approximation of a function around its extremum where linear terms in *x* and *y* vanish. Next, consider a linear approximation of the function *f*(*t*): *f*(*t*) = *f*(*t*_0_) + *f*’(*t*_0_)(*t* – *t*_0_). We look for linear functions

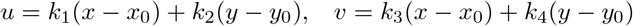

satisfying the equality

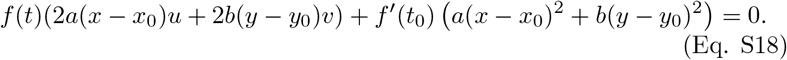

Then

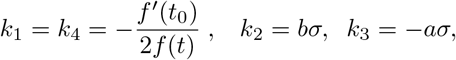

where *σ* is an arbitrary real number which should be determined from the boundary conditions. Thus, linear approximation of the flow field can be determined up to one arbitrary constant.

Since *u*(*x*_0_,*y*_0_) = *ν*(*x*_0_,*y*_0_) = 0, then it is a singular point of the flow field. In order to determine its type, consider the matrix

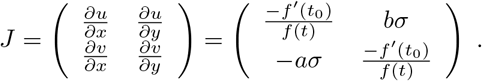

We determine the determinant of this matrix and its eigenvalues:

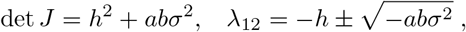

where 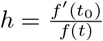. Depending on parameters, the eigenvalues of the matrix *J* can be as follows.

- If *h* > 0 and *ab* > 0, *σ* ≠ 0 then the eigenvalues have negative real parts and nonzero imaginary parts. The corresponding singular point is a stable focus. If *ab* = 0 or *σ* = 0, then it is a stable node with equal eigenvalues. Uncertainty in the choice of *σ* can change focus to node, but in both cases they are stable. The winding number (or index of stationary point) equals 1.
- If *h* < 0 and *ab* > 0, then the singular point is unstable focus or node. The winding number equals 1.
- If *ab* < 0 and *σ* ≠ 0, then there are two different cases depending on the sign of the inequality 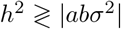. In the case of upper inequality, both eigenvalues have the same sign, and the single point is stable or unstable node. If the inequality is opposite, then the eigenvalues have opposite signs, and the singular point is saddle. Since we expect to have a saddle in the case of opposite signs of the constants *a* and *b*, then |*h*| should be sufficiently small, that is, |*f*’(*t*_0_)| is small enough. If this is not the case, then the linear approximation does not give correct result. The choice of *σ* does not change the type of the singular point provided that condition on *h* is satisfied.

Let us note that Laplacians vanish on linear functions *u* and *ν*. Therefore we obtain an exact solution of equations (Eq. S14), (Eq. S15).

### Travelling waves

Consider a particular form of quadratic function

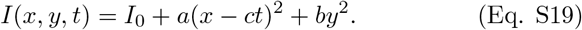

Then

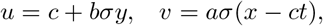

where *σ* is an arbitrary real number. If we suppose that the flow field at the boundary does not depend on time, then *σ* = 0. Hence, *u* = *c,v* = 0, and the flow field does not have singular points.

### Intuitive considerations about the flow field

All examples considered above can be summarized in the following way. Consider level lines of the function *I*(*x,y,t*) on the plane (*x,y*) in two close moments of time, *t* = *t*_0_ and *t* = *t*_1_ (Figure S7).

**Figure S7.**
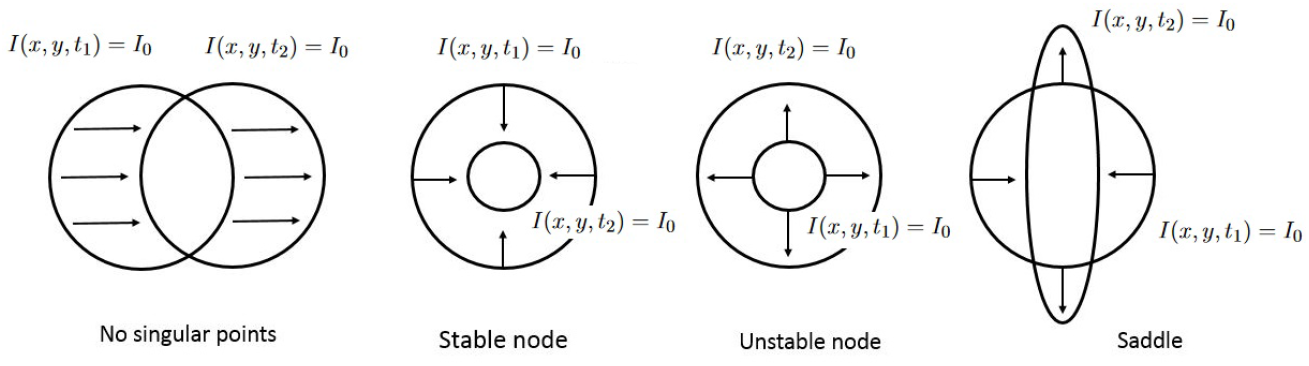
Schematic representation of different time dynamics of the function *I*(*x,y,t*) without singular points (left), with stable or unstable nodes (middle), and with saddle (right).

If the second level line is obtained from the first one by translation (winding number 0), then there are no singular points. If the second level line is obtained by retraction and it is inside the first one (winding number 1), then it is a stable node or a stable focus. In the opposite case, it is unstable node or focus (winding number 1). Finally, if it is partially expanded and partially retracted, then it is saddle point (winding number — 1). Let us recall that winding numbers can be obtained here as *signum* of the product of the eigenvalues.

We will use this empirical definition of singular points to characterize some other types of functions for which explicit analytical solution cannot be constructed.**Standing waves.** Consider the function

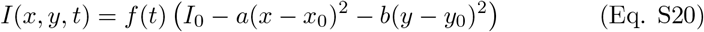

which can be considered as a model of standing wave. As before, we consider a linear approximation of its time dependence

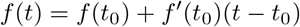

and suppose that *a,b* > 0. According to the previous paragraph, consider level lines of the function *I*(*x, y, t*) around its maximum at *x* = *x*_0_, *y* = *y*_0_ assuming for certainty that *f* (*t*_0_) = 1, *f*’(*t*_0_) > 0. Then *I*(*x*_0_, *y*_0_, *t*_0_) = *I*_0_, *I*(*x*_0_, *y*_0_, *t*_1_) = *I*_1_ > *I*_0_ if *t*_1_ > *t*_0_. Consider the level lines *L*_0_ and *L*_1_ determined, respectively, by the equations

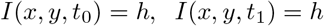

for some *h* < *I*_0_. Both of them are circles, and *L*_0_ is located inside *L*_1_. According to the considerations in the previous paragraph, (*x*_0_, *y*_0_) is unstable node (source).

If *h* approaches *L*_0_, the ratio of circle radii increases and tends to infinity. Therefore, we can expect that the corresponding flow velocity also tends to infinity in the vicinity of the singular point.

Together with this geometrical approach consider equations (Eq. S14), (Eq. S15) with *α* = 0. The flow field *u, v* should satisfy the equality

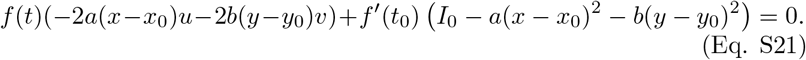

We set

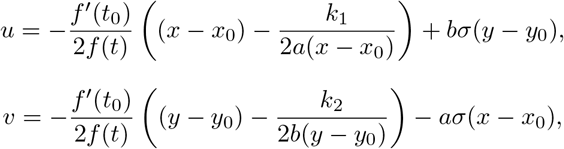

where *σ* is an arbitrary real number, *k*_1_, *k*_2_ are positive and such that *k*_1_ + *k*_2_ = *I*_0_. In agreement with the geometrical considerations, the leading order terms in the flow velocity determine unstable node and the flow velocity tends to infinity at the singular point. This flow field satisfies equations (Eq. S14), (Eq. S15) for *α* = 0. Small positive *α* provides regularization of solution.

## Footnotes

1 For numerical reasons, there would be variation at the very beginning stage, after the pipeline became stable.

2 For a brief introduction of projection problems and of the different interpolation methods that have been used with EEG data (spline surfaces, 2D projection), see [42]: https://www.egi.com/images/kb/SplineInterpolation.pdf

